# Morphological Differences in Isolated Brook Trout Populations in the Desolation Wilderness of the Sierra Nevada

**DOI:** 10.1101/2022.08.15.503945

**Authors:** Lukas Luby-Prikot, Oliver Bock, Joseph Martin

## Abstract

Brook trout (*Salvelinus fontinalis*) is an invasive species in the Desolation Wilderness of California. But it is unknown to what extent this species is evolving to adapt to isolated high altitude lakes. We quantified morphological differences between three brook trout populations in Desolation Wilderness that are in isolation and of common origin. We took standardized photos of fish, created geometric nets of each specimen using points located at known morphological features, and performed a Procrustes superimposition and principal component analysis to examine and cluster morphological variation between individuals. Together, our results show morphological differences between three *Salvelinus fontinalis* populations in independent environments. Our results suggest that invasive species introduced from one source can show physical variation generations after introduction, and thus deserve attention for adapting to and perhaps becoming an increasingly complex part of their ecosystem.

## Introduction

Non-native trout are considered a threat to native species in the mountains of the American West because they predate on endangered amphibians (1)(2) and displace native trout (3), and much attention is paid to how they shape ecosystems. But how invasive trout species are evolving after being introduced to local environments remains an open question.

Brook trout were first introduced to the Grouse, Hemlock, and Upper Doris Lakes of Desolation Wilderness, which are presumed to have been formerly fishless, in the 20th century as a part of fisheries management efforts to provide a recreational fishery. Hemlock Lake was stocked with fish between 1935 and 2000, Grouse Lake was stocked between 1934 and 1985, and Upper Doris was stocked between 1937 and 1942 (4). Hemlock Lake was stocked with 37,040 brook trout from the Tahoe Hatchery and 2,000 from the Mt. Tallac Hatchery (Supp. Note 1), Grouse with 12,000 from Tahoe (Supp. Note 2), and Doris with 22,000 from Tahoe (Supp. Note 3). Stocking stopped at the turn of the century to assist Sierra Nevada yellow legged frog (*Rana sierrae*) recovery. All three lakes are now managed as self-sustaining fisheries (4). Brook trout mature at two years old and spawn in the fall of every year (5). Assuming a generation time of two years, Grouse, Hemlock, and Upper Doris Lakes have experienced 18, 11, and 40 generations since stocking ended respectively. While the lakes are geographically near to each other, their outlets feature impassible fish barriers that make gene flow improbable, and the watersheds are not connected linearly (4).

Subjectively, the lake environments seem to vary with respect to size, depth, geology, water turbidity, plant communities, and sun exposure. Diversity in body shape is one of the most overt responses members of the same species exhibit in accordance with such differences, and has been found to reflect factors including the presence of predators, habitat, food resources, and life history (6) (7) (8) (9) (10) (11). In fact the lake trout, a cousin species of brook trout, exhibits multiple ecomorphs in response to these pressures within a particular lake environment (12).

We sought to determine whether a significant difference exists in the physical features of the isolated brook trout populations of Grouse, Hemlock, and Upper Doris Lakes. In other words, to what extent have these populations diverged and can we determine if they are morphologically distinct?

## Materials and Methods

### Specimen Capture, Measurements, and Photography

A Pentair Aquatic Ecosystems measuring board (SKU# FMB2) was placed on flat shoreline. An approximately 3 ft tall tripod was positioned next to it in a known configuration such that the iPhone 8 camera would capture the same near-vertical perspective during each trial. Brook trout were captured by use of hook and line (artificial flies), and then placed with wet hands on the measuring board, dorsal fin facing up from the perspective of the camera, and head facing left along the central axis of the board. Their length was recorded from nose to the fork of the caudal fin using the measures on the board, and their photograph taken. 30 fish were captured: ten from Grouse Lake between 11:39 and 20:00 on Jul 23 2021, eleven from Hemlock Lake between 10:15 and 11:32 on Jul 24 2021, and nine from Upper Doris Lake between 14:16 and 15:19 on Jul 25 2021.

### Morphology Nets with Morphologica

Using the program Morphologica (13), brook trout images were marked at each of 17 identified physical landmarks in order to approximate the brook trout’s morphology as a net. These points include (1) snout, (2) corner of mouth, (3) anterior eye, (4) closest point of tangency between outline and eye, (5) closest point of tangency between gills and outline, (6) anterior dorsal, (7) posterior dorsal, (8) anterior adipose, (9) top of tail fin, (10) where the lateral line terminates, (11) bottom of tail fin, (12) posterior anal fin, (13) anterior anal fin, (14) origin of pelvic fin, (15) lowest point of body curve, (16) origin of pectoral fin, (17) posterior jaw. Morphologica converted these points into an ordered list of x,y coordinates, stored in a tab separated values sheet.

### Procrustes Superimposition and PCA analysis with MorphoJ

The tab separated value sheet was inserted into the program MorphoJ (14) as a data set, where the outline of each individual’s “net,” represented by the points in the sheet, was treated as an object. We used MorphoJ to perform a Procrustes Superimposition, which optimally translated, rotated, and uniformly scaled each object in order to achieve a similar placement and size, minimizing the differences in the size and orientation of fish that existed in the original photographs. For each of the nets, we calculated the variance between the average location of the landmarks, and the individual landmarks of each fish. Finally, we performed a principal component analysis on the covariance matrix to condense the variation in the data into vectors.

### Statistical Methods

We performed a single factor ANOVA test on fish length data using the Google Sheets add-on XLMiner (15). We performed a Procrustes ANOVA on the shapes of the optimally scaled, rotated, and translated centroids generated by procrustes superimposition using MorphoJ (14).

## Results

Ten brook trout from Grouse Lake, eleven from Hemlock Lake, and nine from Upper Doris Lake were captured and photographed with a standardized procedure. Fish length was measured from nose to caudal fork (Fig. 1). We found that average fish length did not vary between populations (single factor ANOVA, p=0.503, Fig. 2).

**Fig. 1.**
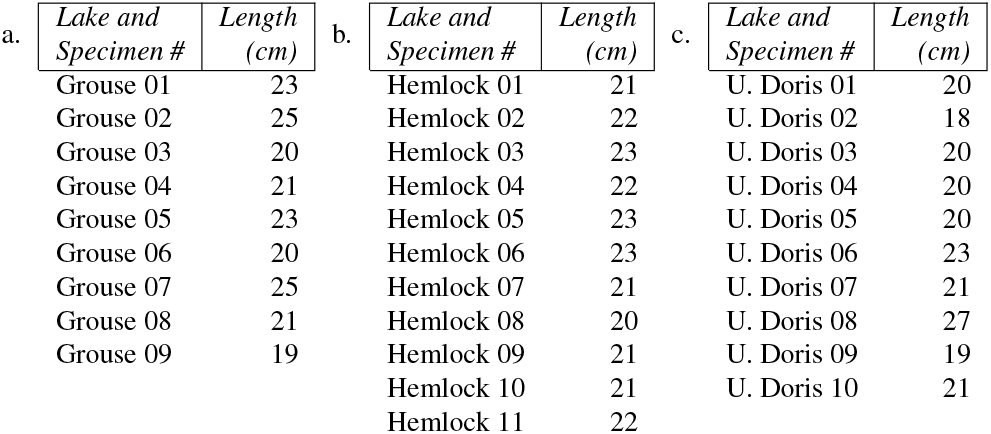
Length of fish caught from Grouse Lake (a.) Hemlock Lake (b.), and Upper Doris Lake (c.). Length was measured from nose to fork of caudal fin.

**Fig. 2.**
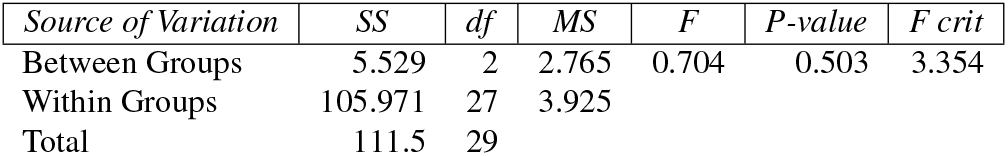
Analysis of variance single factor test using length data from Fig. 1. Displayed are SS (sum of squares), MS (mean square), df (degrees of freedom), F, and P.

Using the program MorphoJ (14), 17 anatomical features were labeled for each fish image (Fig. 3). These points were chosen because they include fixed, non-mobile anatomical features that are similar to ones used in previous bodyshape studies of lake trout (12) and brook trout (16).

**Fig. 3.**
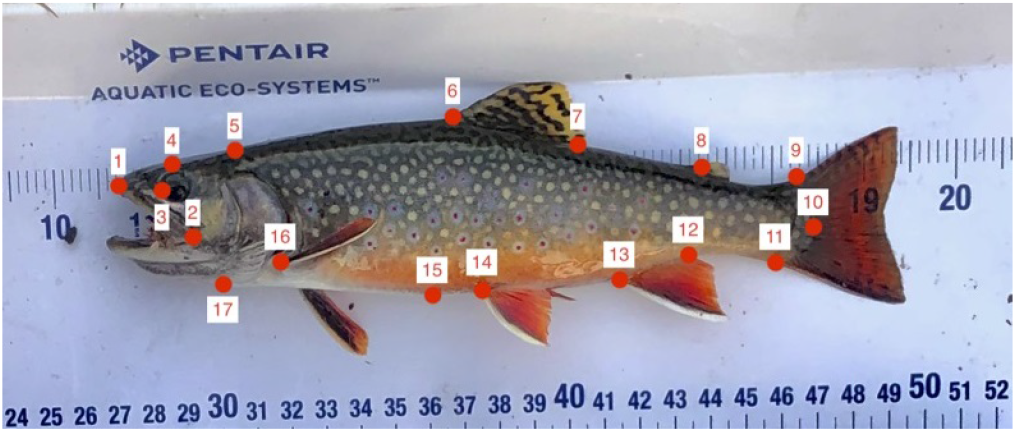
Representative image depicting the orientation of specimens during photography, and the landmarks used to create a mathematical representation of their body shape. Points were pegged to easily identifiable and non moving physical features. These include (1) snout, (2) corner of mouth, (3) anterior eye, (4) closest point of tangency between outline and eye, (5) closest point of tangency between gills and outline, (6) anterior dorsal, (7) posterior dorsal, (8) anterior adipose, (9) top of tail fin, (10) where the lateral line terminates, (11) bottom of tail fin, (12) posterior anal fin, (13) anterior anal fin, (14) origin of pelvic fin, (15) lowest point of body curve, (16) origin of pectoral fin, (17) posterior jaw.

Using the 17 anatomical features, we created geometric nets for each fish, reducing each individual’s morphology to a standardized polygon. A Procrustes superimposition was performed, aligning the points for subsequent PCA analysis (Fig. 4).

**Fig. 4.**
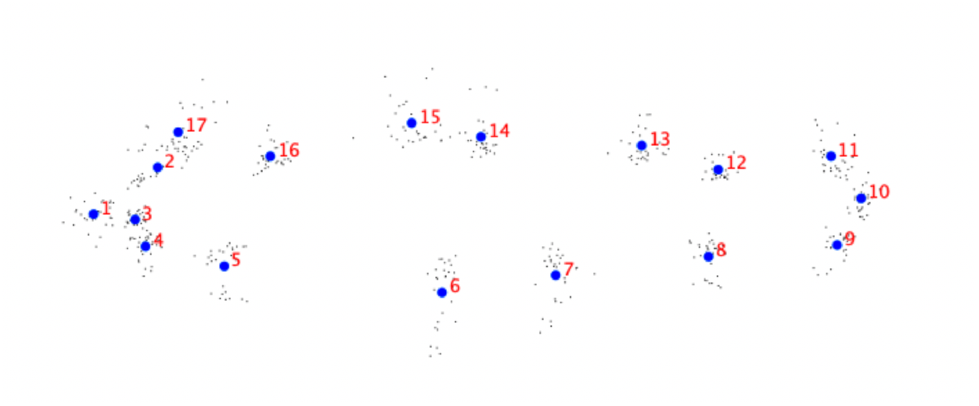
The morphological nets after Procrustes superimposition. Blue dots labeled 1-17 indicate the average position of each known morphological point. Grey dots represent the position of points from individual nets.

A covariance matrix was produced and a principal component analysis was done on the covariance matrix. The first principal component explained 56.1 percent of the variance, the second 13.9 percent of the variance, the third 8.7 percent, and the fourth 4.9 percent. Together, the first and second principal component explain 70 percent of the variance in the data (Fig. 5).

**Fig. 5.**
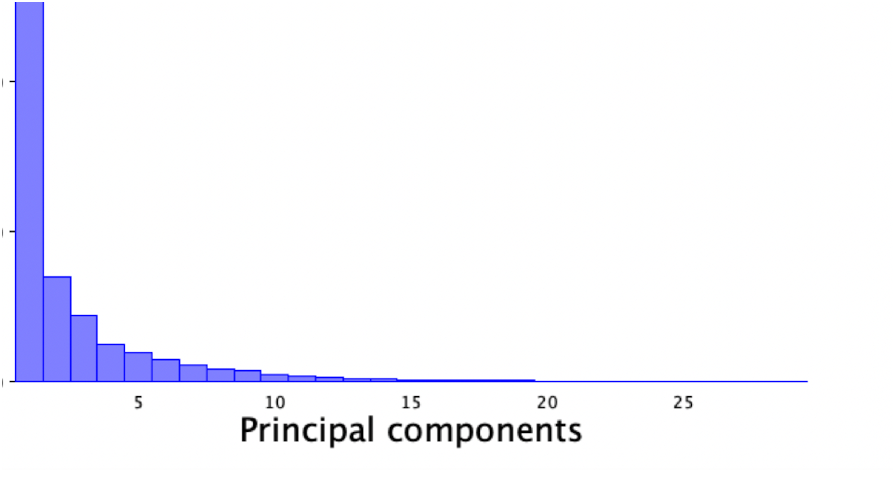
The percent variance in the data represented by each principal component across all samples from all three lakes.

The first principal component was associated most strongly with vectors in shape space that seem to indicate body ‘height’. The greater an individual’s score with respect to the first PC, the more its body was deformed vertically, especially from landmark 17-13 on the top, and 5-8 on the bottom (Fig. 6).

**Fig. 6.**
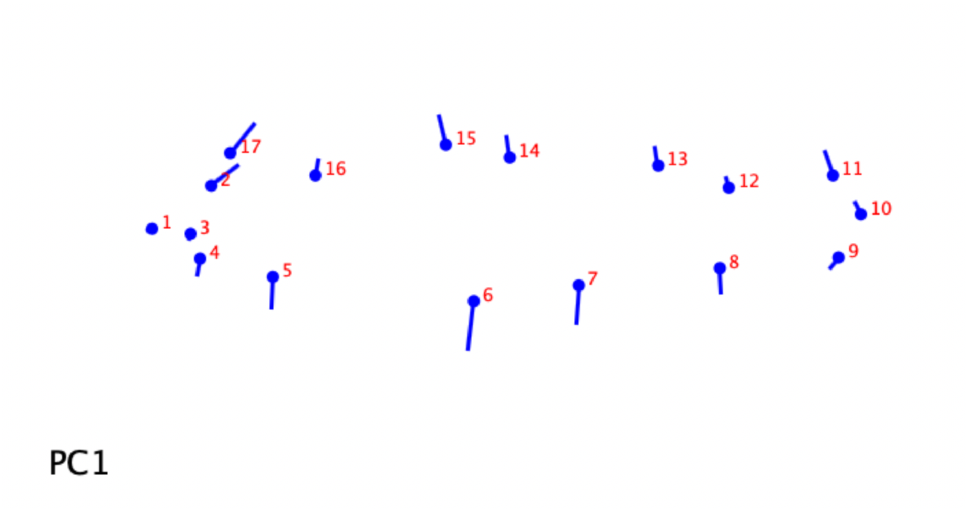
Lollipop diagram depicting the vectors and relative magnitudes of deviations from the mean shape associated with the first principal component.

**Fig. 7.**
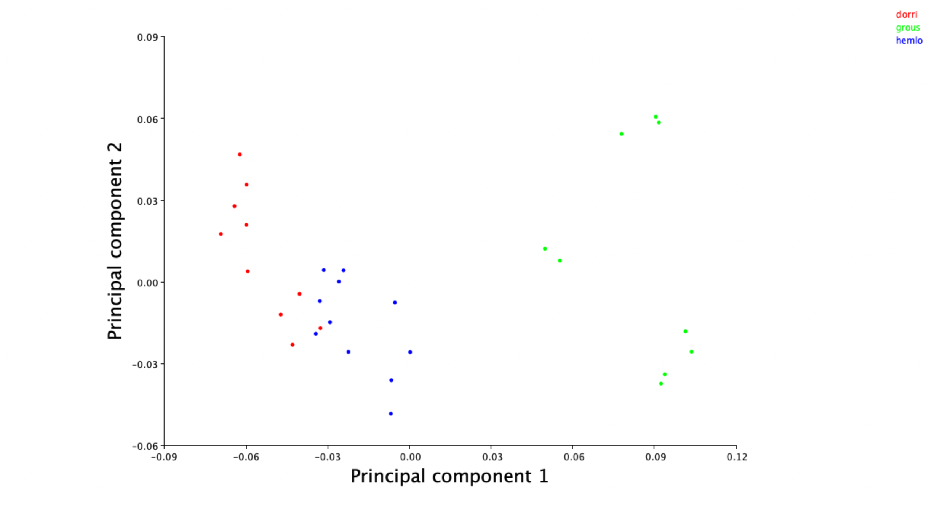
Scatter plot relating the first and second principal components of morpho-logical variance in fish sampled from Upper Doris (red dots), Grouse (green dots), and Hemlock Lake (blue dots). Each dot represents the first and second PCA value for an individual fish.

Using PC 1 and PC 2, which contained the majority of the variation in the data, as axes for a scatter plot revealed that brook trout body morphology clusters by population around distinct centers. Visually, the edges of those clusters differ between Grouse Lake and the other two lakes, while Doris and Hemlock exhibit some overlap (Fig. 8).

**Fig. 8.**
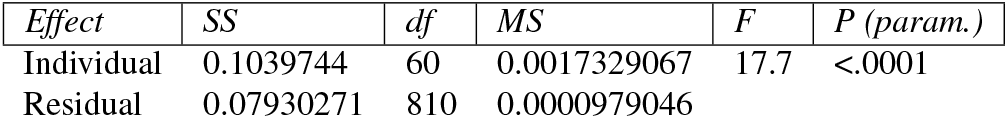
Procrustes analysis of variance test performed on morphological shape data using lake origin as a classifier. Displayed are SS (sum of squares), MS (mean square), df (degrees of freedom), F, and P.

We performed a Procrustes ANOVA on the coordinates of each landmark after being translated, rotated and scaled by Procrustes superimposition. We used “lake” as a classifier to partition variance in shape between the three populations. The results of our Procrustes ANOVA show that there was a statistically significant difference in body shape between the three populations (p<0.0001, Fig. 8).

## Discussion

Together, the results of the principal component analysis and Procrustes ANOVA show that brook trout body shape morphology is different between the three lake basins. This has implications for how brook trout may be functioning within their ecosystems. In a previous study, hatchery brook trout have been shown to change genetically and phenotypically in response to their environments, displaying distinct morphology between different populations (16). Our results, corollary to these, call attention to invasive brook trout populations across the American West for perhaps achieving the same.

Our methodology for assessing morphology comes with distinct, novel advantages as a tool for research and population management. It relies on simple optical methods for which the tools are everyday items or easily procured, and low in volume and weight. It does not require anaesthetization or the taking of perishable specimens, making it ideal for surveying remote, small populations. In spite of these things, it retains the statistical power necessary to make inferences about population morphology with a limited number of samples. It is easy to execute, and relies on publicly accessible programs (13)(14).

We do not know the exact causes of the difference of morphology we observed. One explanation is natural selection driven by differences in the lake environments causing changes in allele frequencies of morphologically relevant genes. It is also possible that the variation between populations was caused by a founder effect, although this seems less likely due to the continuous stocking from California state hatcheries. Previous studies have used genetic analysis of brook trout mitochondrial and nuclear DNA sequences to establish differences between populations (17)(18)(19)(20). We have begun efforts to establish the genetic similarities of these populations through sequencing of the mitochondrial DNA gene *NADH4*, but it will be challenging to establish a causal relationship between genetic differences and morphological changes.

It is also possible the differences in morphology we observed were caused by a developmental or physiological response to environmental pressures such as altitude, pH, or temperature. All these factors are known to effect fish morphology (21)(22)(23), so it is possible the differences we observed do not rest with a genetic origin. However, we did find there to be no difference in mean length between the three populations, suggesting that differences in morphology caused by size likely did not play a role. The fact that the variation was concentrated in PC 1, and exhibits itself primarily with respect to body height, seems corollary to the humper ecomorph exhibited by Lake Trout in the Great Lakes region (12). This could be in response to several pressures. The lakes vary in altitude and size, and as indicated in supplementary notes, brook trout were stocked alongside different quantities of other species of fish who could have exerted ecological pressures upon them before dying out.

Unlike the other lakes, Hemlock Lake was stocked with fish from the Mt. Tallac hatchery. However, the last stocking of Tallac fish was the second of twelve stockings in 1936. Stocking would continue at a greater number of brook trout per sortie until 1949. Fish from Tallac may contribute less to the allele frequencies of the Hemlock population. Even if the Hemlock population is disregarded in our analysis, Grouse and Doris Lakes, stocked exclusively from the Tahoe hatchery, cluster even farther apart from each other than from Hemlock on the PC scatter plot. While we have limited information on the genetic origin of the population founders, we believe our method and findings can serve to help manage brook trout populations as they react to a changing climate and other ecological pressures.

## ACKNOWLEDGEMENTS

We thank Mitch Lockhart (California Department of Fish and Wildlife District Biologist, North Central Region) for helping us plan our trip to the Desolation Wilderness, helping us access to stocking records, and volunteering to supervise our research on cite, Sasha Sherstsnev (Massachusetts Institute of Technology) for writing his custom program Morphologica to create mathematical representations of landmarks, and Alexander Luby-Prikot (Boston University) for his working as a lab assistant on the effort to sequence mitochondrial DNA.

---

**Supplementary Note 1:**
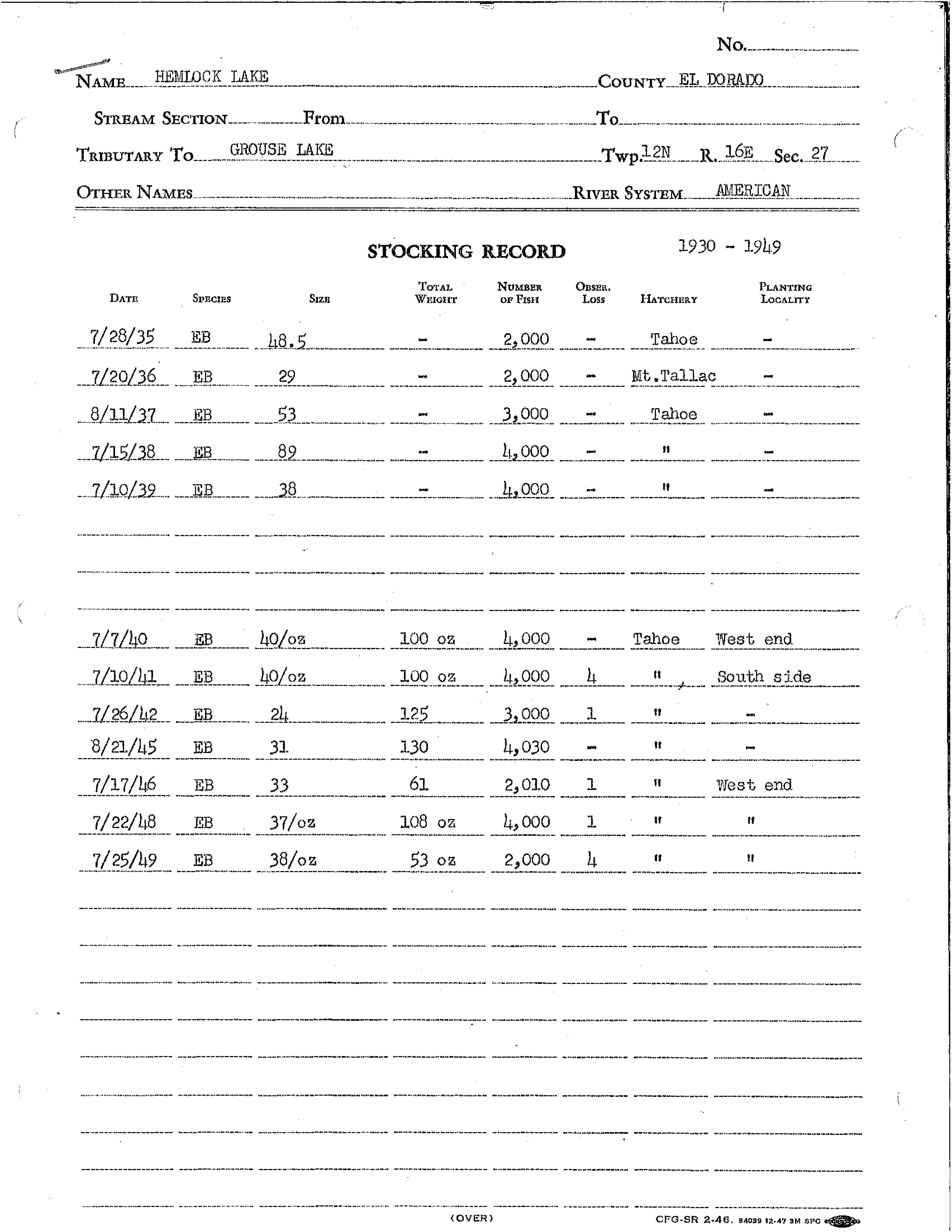
California Department of Fish and Wildlife Brook trout stocking records for Hemlock Lake, 1935-1949.

**Supplementary Note 2:** California Department of Fish and Wildlife Brook trout (EB), Rainbow trout (RT), and Steelhead (SH) stocking records for Grouse Lake, 1930-1949.

**STOCKING RECORD 1930 - 1949**
| DATE | SPECIES | SIZE | TOTAL WEIGHT | NUMBER OF FISH | OBSER. LOSS | HATCHERY | PLANTING LOCALITY |
| --- | --- | --- | --- | --- | --- | --- | --- |
| 7/1/30 | RT | - | - | 5,000 | - | Mt. Tallac |  |
| 7/24/33 | SH | - | - | 4,000 | - | " |  |
| 7/15/34 | EB | 53.5 | - | 4,000 | - | Tahoe |  |
| 7/20/36 | RT | 35 | - | 3,000 | - | Mt. Tallac |  |

NOTE: For additional possible planting records see: Unidentified stocking records El Dorado County.

|  |  |  |  |  |  |  |  |
| --- | --- | --- | --- | --- | --- | --- | --- |
| 7/7/40 | EB | 40/oz | 75 | 3,000 | - | Tahoe | - |
| 7/10/41 | EB | 40/oz | 75 oz | 3,000 | - | " | North side |
| 7/26/42 | EB | 24 | 84 | 2,000 | 1 | " | - |
| 7/10/44 | RT | 21 | 172 | 3,610 | 3 | " | Upper end |
| 8/19/45 | RT | 15 | 400 | 6,000 | 3 | " | - |
| 7/17/46 | RT | 21 | 238 | 5,000 | 2 | " | South shore |
| 7/18/47 | RT | 23/oz | 261 oz | 6,000 | 5 | " | North side |
| 7/22/48 | RT | 28/oz | 215 oz | 6,020 | 5 | " | " |
| 7/25/49 | RT | 36/oz | 167 oz | 6,000 | 35 | " | " |

(OVER)
CFG-SR 2-46, 43009 4-51 3M © SPO
STREAM SECTION From To

**Supplementary Note 3:**
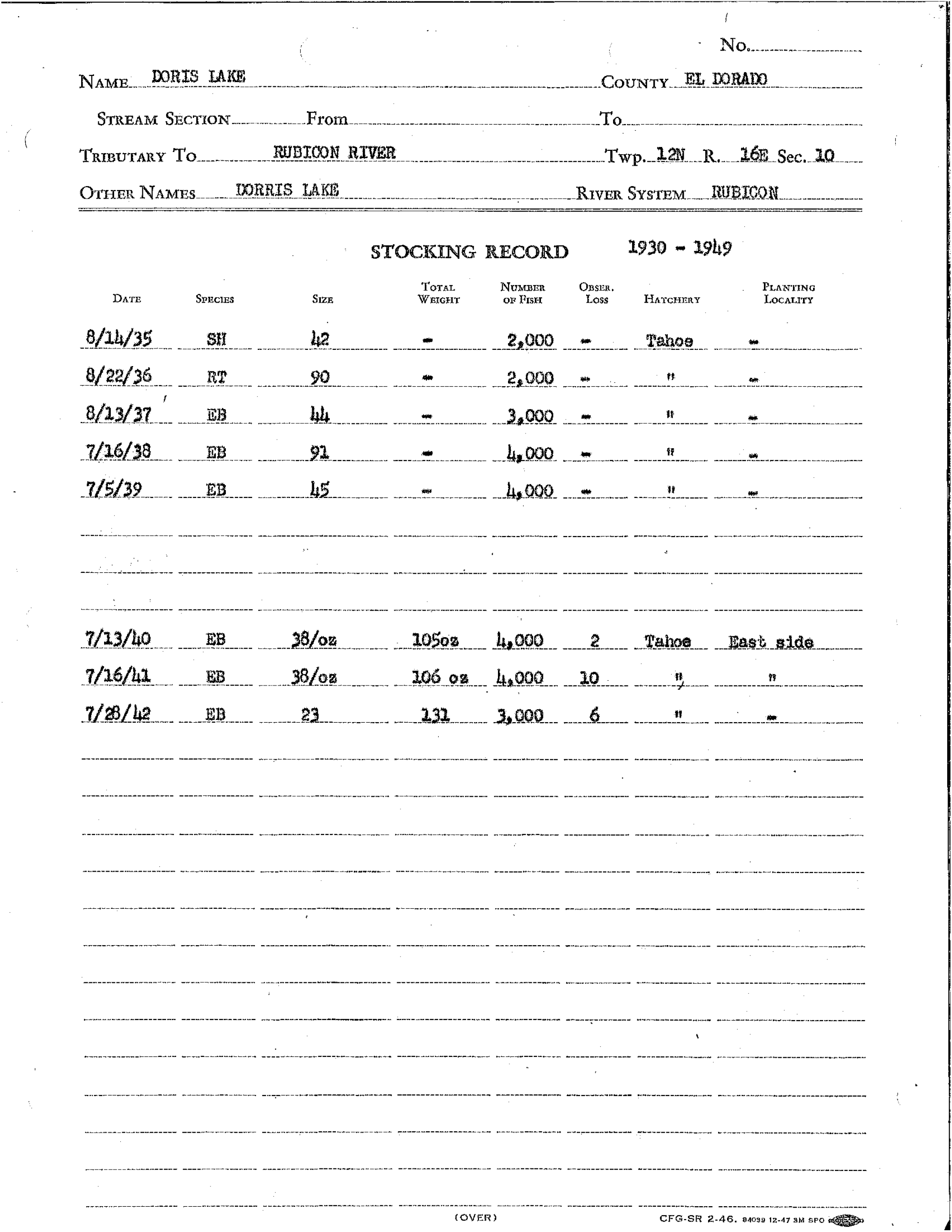
California Department of Fish and Wildlife Brook trout (EB), Rainbow trout (RT), and Steelhead (SH) stocking records for Doris Lake, 1935-1942.

